# DNA Methyltransferase 1 and 3a Expression in the Frontal Cortex Regulates Palatable Food Consumption

**DOI:** 10.1101/2021.05.23.445176

**Authors:** Mohan C Manjegowda, Jonathan Joy-Gaba, Eric Wengert, Anusha U. Saga, Daniel Warthen, Amelie Kuchler, Ronald Gaykema, Manoj K. Patel, Nathan C. Sheffield, Michael M. Scott

## Abstract

DNA methylation is an important regulatory mechanism in the control of neuronal function. Both during development and following exposure to salient stimuli, plasticity in the methylation of cytosine residues leads to a change in neuron excitability that subsequently sculpts animal behavior. However, although the response of DNA methyltransferase enzymes in adult neurons to stimuli such as drugs of abuse have been described, less is known about how these enzymes regulate methylation at specific loci to change the drive to ingest natural rewards. Specifically, we do not understand how changes in methylation within important brain areas known to regulate palatable food intake can affect ingestion, while a detailed investigation of the neurophysiological and genomic effects of perturbing methyltransferase function has not been pursued. By deleting DNA methyltransferase 1 and 3a in the mouse prefrontal cortex, we observed the requirement for these enzymes in the regulation of nutrient rich food consumption in the absence of any effect on the intake of low fat and low sugar chow. We also determined that the deletion profoundly affected neuron excitability within pyramidal cells resident in superficial layers II/III of the cortex but had little effect in deep layer V neurons. Finally, reduced representation bisulfite sequencing revealed both hypo and hypermethylation in response to methyltransferase deletion, an effect that was observed in binding sites for retinoic acid receptor beta (RARβ) located within regulatory regions of genes known to affect neuronal function. Together, our data suggest that alterations in the actions of RARβ could shift neuronal activity to reduce palatable food intake.

## Introduction

DNA methylation is one of the most extensively studied epigenetic marks and is implicated in many biological processes such as neuronal development, X-chromosome inactivation and imprinting. Yet, it has only recently been appreciated that DNA methylation also has an active role in the dynamic regulation of transcription in terminally differentiated cells(Guo et al., 2011; Mo et al., 2015; Patil et al., 2014; Pinney, 2014; Varley et al., 2013). While several DNA methyltransferase (DNMT) isoforms were shown to be expressed in terminally differentiated neurons(Feng et al., 2005; Feng et al., 2010; Goto et al., 1994; Inano et al., 2000; Kadriu et al., 2012), discovery of demethylating enzymes belonging to the TET (ten eleven translocation) family of proteins, which actively catalyze DNA demethylation, further supports the presence of bidirectional control over DNA methylation in mature cells (Ginno et al., 2020; Pastor et al., 2013; Yu et al., 2015). Subsequent studies have shown how DNA methylation writers (DNMT) and erasers (TET and TDG) are known to be interdependent in terms of their activity and spatial recruitment, depending on context (Gu et al., 2018), with the balance of methylation and demethylation determining the methylation level for a given locus (Ginno et al., 2020). An understanding of where this dynamic regulation of DNA methylation may occur in the mature neurons has also evolved significantly. Methylation of cytosine in non-CpG contexts, for example, was initially believed to act as a mark restricted to developmentally active cells(Pinney, 2014). However, with the advent of genome-level high throughput techniques, although relatively less frequent compared to levels seen in pluripotent cells, an increasing number of studies have identified non-CpG methylation in mature brain tissue(Mo et al., 2015; Pinney, 2014; Varley et al., 2013).

At the level of animal behavior, plasticity within patterns of DNA methylation has been shown to be crucial following exposure to salient stimuli (Feng et al., 2010). Within adult animals, the actions of two DNMT isoforms that mediate the initial addition of the methyl group in mature neurons, DNMT1 and DNMT3a (Feng et al., 2005; Inano et al., 2000; Kadriu et al., 2012), are required for the development of synaptic plasticity during spatial learning tasks and fear conditioning (Miller and Sweatt, 2007).

Reward-driven learning (LaPlant et al., 2010) and ingestive behavior (Kohno et al., 2014) have also been shown to be regulated by DNA methylation, suggesting that DNA methylation may be an important control point acting to modulate consumption. While the DNMT enzymes have been shown to be important regulators of drug reward seeking behavior (LaPlant et al., 2010; Massart et al., 2015; Wright et al., 2015), conflicting data published in separate studies demonstrate the complexities of this biological process. Although restricting DNMT3a activity enhanced the development of a conditioned place preference towards cocaine (LaPlant et al., 2010), pharmacological inhibition of the DNMT enzymes resulted in an impairment in cued cocaine seeking (Massart et al., 2015), with whole animal methionine supplementation also reducing reinstatement of responding for this reward (Wright et al., 2015).

Unfortunately, although there is still much to be learned regarding the role of DNA methylation in regulating cocaine reward driven behavior, there is even less known about how natural reward intake is regulated by this epigenetic process. Furthermore, given that prior work has shown how whole animal shifts in methylation can produce significantly different effects on cocaine ingestion when compared to sucrose consumption (Wright et al., 2015), it is quite likely that the epigenetic changes that result in altered pursuit of these rewards also differ, necessitating the investigation of DNMT action specifically within this context.

Only one study has thus far investigated the role of DNMT enzyme activity on feeding behavior, with prior work showing how DNMT3a, but not the second isoform highly expressed in adults, DNMT1, is required in the paraventricular nucleus of the hypothalamus for the regulation of chow food intake, body weight and adiposity (Kohno et al., 2014). Interestingly, loss of function of the methyl cytosine binding protein, MECP2, has shown that impaired readout of genomic methylation may also affect palatable food intake, possibly explaining the increased incidence of obesity and food intake in patients with Rhett syndrome (Fukuhara et al., 2019). Currently, however, little is known about how or whether manipulation of DNMT expression could regulate palatable food intake and how changes in DNMT expression affects locus-specific methylation patterns genome wide.

To investigate how DNA methylation may regulate palatable food intake, we chose to study the frontal cortex, as this brain area is involved in the control of food reward consumption, as opioid signaling in the prelimbic and infralimbic regions has been shown to bidirectionally drive the intake of high calorie food (Blasio et al., 2014; Castro and Berridge, 2017; Mena et al., 2011). We deleted two DNMT isoforms expressed in the adult cortex, DNMT1 and DNMT3a, and discovered that this significantly reduced palatable food consumption, but had no effect on the intake of low fat and low sugar chow. While we were able to confirm that DNMT expression within the mPFC is required for trace fear conditioning (Morris et al., 2014), our manipulation did not change affect or stress responsiveness. With respect to mechanism of action, our studies revealed that the observed reduction in food intake was accompanied by significant variation in the effect of DNMT deletion on neuronal excitability within pyramidal cell layers of the frontal cortex. Finally, our analysis of the DNA methylation pattern changes that occurred with DNMT1 and DNMT3a deletion showed a surprising effect within consensus binding sites for retinoic acid receptor beta (RARβ). Together, our data suggest that alterations in the actions of RARβ could be producing a shift in neuronal activity that results in a reduction in palatable food intake.

## Results

### Deleting *DNMT1* and *DNMT3a* in medial prefrontal cortex neurons and isolation of a genetically defined population of DNMT-null pyramidal neurons

Aberrant DNA methylation within the prefrontal cortex (PFC) has been observed across pathological conditions like depression, stress, addiction, eating disorders and neurodegenerative disorders. DNA methylation has been implicated in the development of synaptic plasticity and in the control of intrinsic membrane properties (Feng et al., 2010; Levenson et al., 2006; Meadows et al., 2015; Miller et al., 2008). These studies have used either pharmacological intervention or conditional knockout of DNMT enzymes to examine effects transcriptional, cellular, and behavioral changes, almost exclusively in the hippocampus or throughout the forebrain. Functional consequences of DNA methylation alterations made selectively in the PFC are, consequently, not completely understood. Furthermore, genome-wide, the molecular details as to how DNA methylation operates to maintain neuron function are not clear. To this end, we set out to assess how deletion of the primary adult-expressed methyltransferases, *DNMT1* and *DNMT3a*, from the frontal cortex, affects behavior in addition to CpG and non-CpG methylation levels. Here, we spatiotemporally deleted the enzyme active site from either *Dnmt1* or *Dnmt3a* (SKO), or both simultaneously (*DNMT1/3a* double knockout, DKO) in the PFC using a neuron-specific promoter-driven Cre recombinase (Figure 1A). Following AAV2.1-Syn-Cre-GFP or control AAV2.1-syn-GFP injection into *DNMT1* and *DNMT3a* double-floxed mice, we confirmed targeting to the PFC with detection of GFP expression in coronal sections of PFC (Figure 1B). Within the PFC, GFP expression was observed in regions extending dorsally, including the cingulate cortex and prelimbic region, and regions extending ventrally, including the infralimbic region. Our injections also extended to caudal portion of the orbitofrontal cortex. To determine the extent of deletion following Cre recombinase expression, we used a flow-cytometry-based sorting strategy (Figure 1C, see methods) to isolate a genetically defined population of PFC pyramidal neurons (PNs), then subsequently measured mRNA expression levels of *DNMT1* and *DNMT3a* by qRT-PCR. Using SATB2, a marker specific to cortical excitatory neurons (Gaykema et al., 2014), we sorted the PNs from the PFC tissue punches and confirmed that mRNA levels of both *DNMT1* and *DNMT3a* were significantly reduced in GFP and SATB2 double positive nuclei from DKO mice as compared to GFP-negative SATB2-positive nuclei isolated from the same animal (Figure 1D). Collectively, these results establish the successful deletion of *DNMT1/3a* within the PFC.

**Figure 1.**
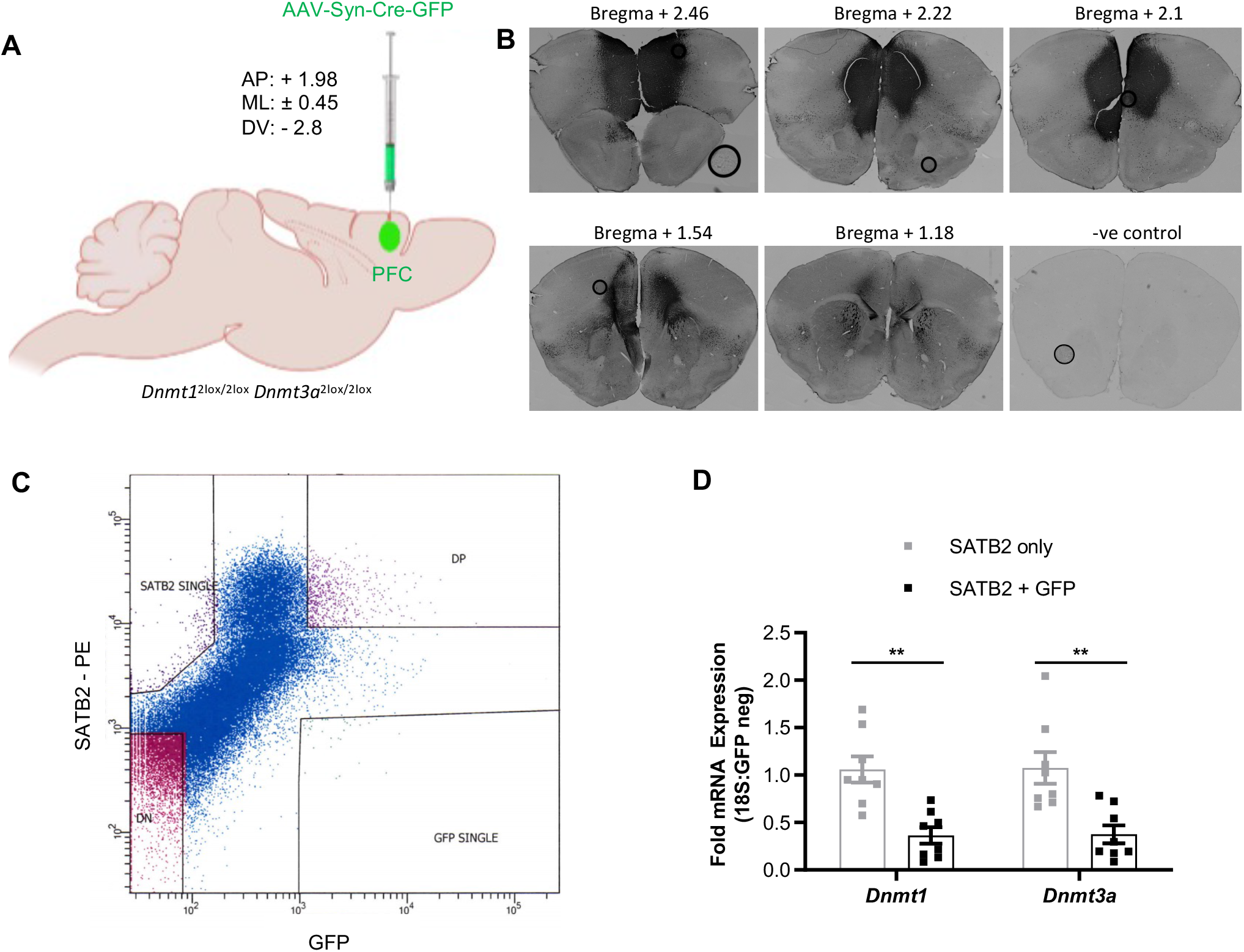
Neuron-specific deletion of DNMT1 and 3a in the PFC. A. Schematic diagram showing injection details for conditional knockout of DNMT1 and DNMT3A in mPFC neurons. B. Representative images of PFC coronal sections showing the rostral/caudal and dorsal/ventral expression of Cre-GFP. C. A representative scatter plot from flow cytometer showing the distribution of SATB2 and/or GFP positive nuclei isolated form the mPFC tissue punches of synapsin-Cre-GFP virus injected DNMT12lox/2lox DNMT3a2lox/2lox mice. D. Relative mRNA expression levels of DNMT1 and DNMT3A in FACS-sorted SATB2-only positive (GFP-negative) and SATB2 and GFP doublepositive neuronal nuclei confirming DNMT deletion (n=8, paired t-test **p < 0.01). Data information: mean ± SEM

### *DNMT1 and DNMT3a* deletion in the PFC significantly reduces palatable food intake

Recent work from our group as well as others has implicated PFC in the control of feeding and food-driven exploratory behavior (Gaykema et al., 2014; Mena et al., 2011; Sinclair et al., 2019; Warthen et al., 2016), potentially acting to selectively regulate palatable food consumption.

Using a paradigm we previously employed to investigate binge-like food intake (Gaykema et al., 2014), we assessed whether conditional knockout of one or both DNMT enzymes had any effect on the consumption of palatable food. Surprisingly, DKO mice consumed significantly less Western Diet (WD, 4.41kcal/g, Envigo TD.88137), containing high levels of sucrose and lipids, than did controls (Control, n=12, 0.998 ± 0.04g; DKO, n=12, 0.858 ± 0.054g) (Figure 2A). We then repeated the binge-feeding assay with a separate cohort of PFC DKO mice to assess whether suppressed consumption was dependent upon the composition of the palatable food. Using HFD (5.56 kcal/g, D12331, Research Diets) a diet where the percentage of fat kcal per gram is significantly greater than that of sucrose (unlike the balanced distribution of sucrose and fat in WD), we again observed significantly less HFD consumption by PFC DKO mice (n=12, 0.363 ± 0.039g) compared to controls (n=12, 0.591 ± 0.065g) (Figure 2A). However, normal chow consumption was not altered by the PFC deletion of the DNMT enzymes(control, n=12, 0.157 ± 0.021g; DKO, n=12, 0.130 ± 0.021g) (Figure 2A). We then tested whether the suppression of palatable food intake could be maintained following repeated exposure to either WD or HFD, to examine whether diet novelty was required for the observed effect of DNMT deletion on feeding. Interestingly, PFC DKO mice showed suppression of intake on HFD only on the first day of binge feeding (Control, n=12, Day 1, 0.590 ± 0.065g, Day 2, 0.641 ± 0.044g, Day 3, 0.686 ± 0.054g; DKO, n=12, Day 1, 0.362 ± 0.038g, Day 2, 0.545 ± 0.050g, Day 3, 0.589 ± 0.052g Figure 2B). However, PFC DKO mice fed WD showed suppressed binging activity on all three days of testing (n=12, Day 1, 0.858 ± 0.054g; Day 2, 0.737 ± 0.063g; Day 3, 0.833 ± 0.075g) when compared to controls (n=12, Day 1, 0.998 ± 0.040g; Day 2, 0.997 ± 0.059g; Day 3, 1.077 ± 0.065g) on all three days (Figures 2C). We then investigated whether the effect could be recapitulated by single conditional knockouts of either *DNMT1* or *DNMT3a* in PFC. However, SKOs did not alter the binge-like feeding on WD (*DNMT1* SKO Control, n=10, 0.817 ± 0.043g; *DNMT1* SKO, n=12, 0.774 ± 0.038g; *DNMT3α* SKO Control, n=20, 0.644 ± 0.056g; *DNMT3α* SKO, n=14, 0.575 ± 0.047g) (Figure 2D).

**Figure 2:**
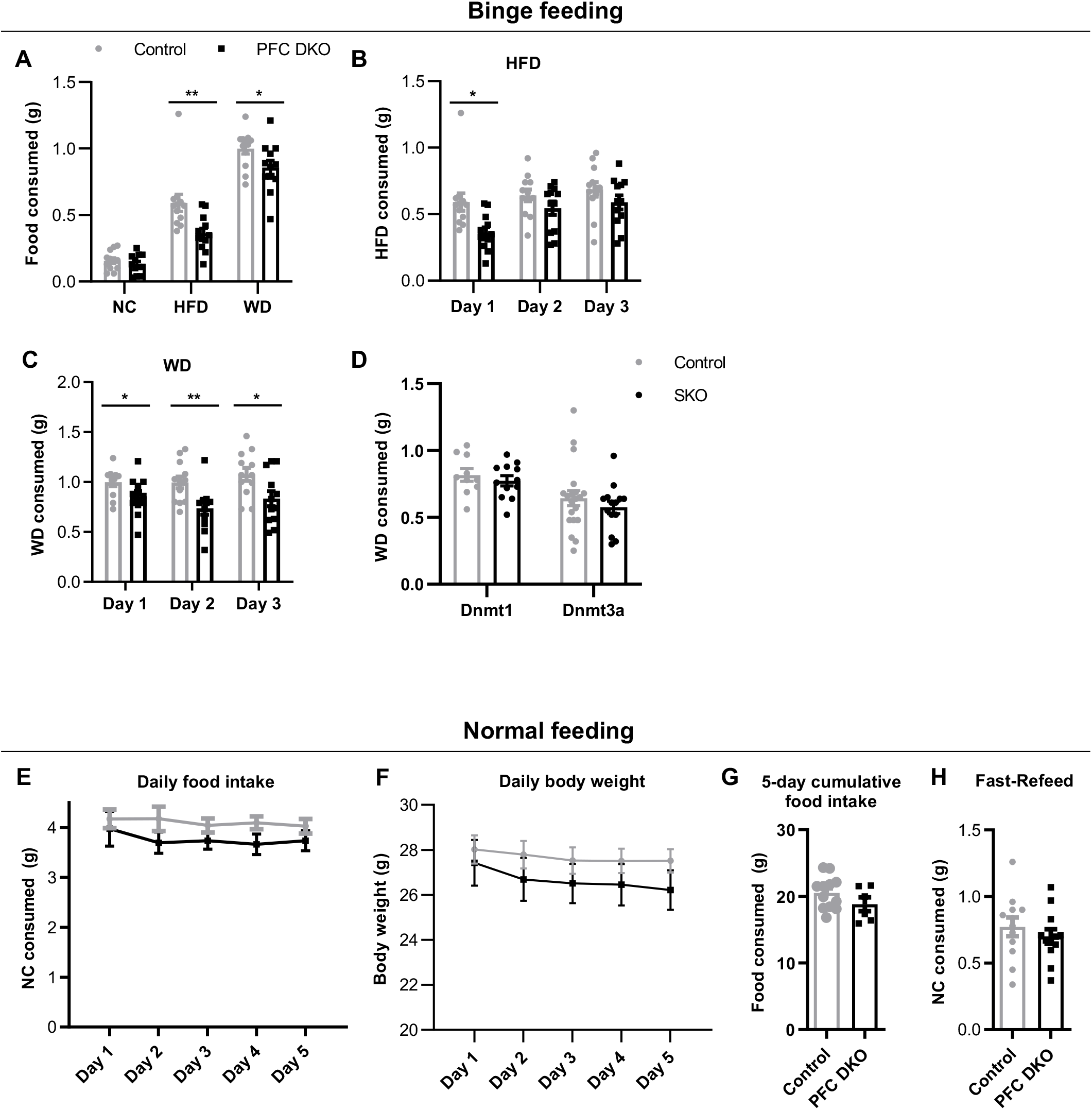
Deletion of *DNMT1* and *DNMT3a* in the mPFC reduces palatable food intake without altering normal feeding behavior. A. Consumption of various diets by control and PFC-DKO groups as measured by a binge-like feeding assay. PFC-DKO group showed suppression of binge feeding on high fat diet (HFD) and western diet (WD) but not on normal chow (NC). (n=12, Student’s *p*-test \**p*<0.05, \*\**p*<0.01). B. Consumption of HFD by control and PFC-DKO groups as measured by a binge-like feeding assay. Knockout-mediated binge feeding suppression is attenuated over time. (n=12, 2way ANOVA, Post-Hoc Bonferroni multiple comparisons \**p*<0.05). C. Consumption of WD by control and PFC-DKO groups as measured by a binge-like feeding assay. The knockout-driven reduction in binge feeding is persistent over time. (n=12, 2way ANOVA, Post-Hoc Bonferroni multiple comparisons \**p*<0.05, \*\**p*<0.01). D. Consumption of WD by control and *DNMT1* or *DNMT3A* SKO groups as measured using a binge-like feeding assay. n=10 for *DNMT1*-control, n=12 for *DNMT1*-SKO, n=12 for *DNMT3A*-control and SKO. E. Daily food consumption of NC by control and PFC-DKO groups. No difference in NC consumption (control n=12, PFC-DKO n=6). F. Body weight of control and PFC-DKO groups while monitoring daily food consumption. No difference in body weight was observed (control n=12, PFC-DKO n=6). G. Five days cumulative NC consumed by control and PFC-DKO groups. No difference in food consumption was observed (Control n = 12, PFC-DKO n=6). H. Consumption of NC by control and PFC-DKO groups during a fasting/refeeding paradigm. No difference in NC consumption was observed (n=12). Data information: error bars are ± SEM

Given that consumption of calorically dense, highly palatable food was affected by *DNMT* deletion, we investigated whether *DNMT* deletion could produce a change in normal feeding behavior of rodent chow. Over a period of 5 days, we observed no difference between PFC DKO (n=12, Day 1, 3.978 ± 0.347g; Day 2, 3.697 ± 0.213g; Day 3, 3.737 ± 0.17g; Day 4, 3.665 ± 0.208g; Day 5, 3.738 ± 0.201g) and control (n=6, Day 1, 4.175 ± 0.188g; Day 2, 4.178 ± 0.244g; Day 3, 4.043 ± 0.143g; Day 4, 4.098 ± 0.128g; Day 5, 4.028 ± 0.147g) mice in daily food consumption of normal chow (Figure 2E) or in body weight (Control, n=12, Day 1, 28.0167 ± 0.626g; Day 2, 27.787 ± 0.612g; Day 3, 27.523 ± 0.591g; Day 4, 27.511 ± 0.540g; Day 5, 27.514 ± 0.518g; DKO, n=6, Day 1, 27.425 ± 1.017g; Day 2, 26.687 ± 0.953g; Day 3, 26.51 ± 0.872g; Day 4, 26.45 ± 0.923g; Day 5, 26.217 ± 0.88g) (Figure 2F). Furthermore, no difference in the 5-day cumulative body weight was observed between groups (Control, n=12, 20.523 ± 0.703g; DKO, n=6, 18.815 ±1.046g) (Figure 2G).

It is possible that *DNMT* deletion affects feeding behavior only during periods of increased feeding drive resulting from an increase in food salience, independent of nutrient content. We therefore tested the effect of increasing the salience of rodent chow by subjecting mice to an 18 h overnight fast. Chow was then reintroduced, and consumption was measured for one hour the following morning. Surprisingly, there was no observable difference in regular chow consumption using the fasting/refeeding paradigm (Control, n=12, 0.773 ± 0.071g; DKO, n=12, 0.7 ± 0.056g) (Figure 2H). These results suggest that DNMT enzymes in the PFC selectively regulate palatable food intake that is dependent upon nutrient content. Further, our results also suggest that the functional roles of *DNMT1* and *DNMT3a* in the PFC are overlapping.

### PFC deletion of *DNMT1 and DNMT3a* does not modulate affect or the neuroendocrine response to stress

While prior studies have looked at the effects of *DNMT* deletion throughout the forebrain (Morris et al., 2014; Morris et al., 2016), little work has been done to examine how cortically expressed DNMT enzymes can affect a more limited set of neurons within this brain region. Furthermore, conflicting data make it difficult to assess how DNMT expression could regulate anxiety-like behavior. Initially, we set out to investigate whether our deletion could regulate affect, as prior studies have suggested. Using a sucrose preference test, we observed that both groups showed similar preference for sucrose over water (Control, n=9, H2O, 52.574 ± 7.0%; 0.5% sucrose, 56.457 ± 5.701%; 1% sucrose, 40.328 ± 5.843%; 2% sucrose, 86.503 ± 3.92%; DKO, n=7, H2O, 57.5 ± 7.443%; 0.5% sucrose, 63.808 ± 4.971%; 1% sucrose, 42.071 ± 4.241%; 2% sucrose, 80.745 ± 4.95%) (Figure 3A) indicating that the DKO deletion did not produce anhedonia. Furthermore, DKO animals showed no differences in novel social (Control, n=11, 3.426 ± 0.89; DKO, n=11, 3.377 ± 0.736) (Figure 3B) or novel object (Control, n=12, No Object, 21.585 ± 2.33s; Object, 54.993 ± 13.243s; DKO, n=12, No Object, 21.154 ± 3.342s; Object, 55.714 ± 15.94s) (Figure 3C) interaction when compared to controls nor were any changes in general locomotion (Control, n=8, 3000.660 ± 370.366cm; DKO, n=8, 3624.006 ± 425.77cm) (Figure 3D) observed. As prior work looking at *DNMT1* or *DNMT3a* single knockdown within the PFC had suggested that reduced DNMT expression was either anxiolytic or anxiogenic(Elliott et al., 2016; Morris et al., 2016), we investigated the effect of the DKO on both elevated plus maze performance and locomotion in an open field arena. Surprisingly, PFC DKO mice performed at levels comparable to controls in the elevated plus maze assay (Control, n=8, Closed Arms, 164.306 ± 13.562s; Open Arms, 24.191 ± 7.443s; PFC DKO, n=8, Closed Arms, 157.174 ± 10.536s; Open Arms, 17.209 ± 4.622s) (Figure 3E) as well as in an open field arena (Control, n=8, Center, 31.6 ± 8.505s; Periphery, 268.001 ± 8.376s; PFC DKO, n=8, Center, 33.463 ± 4.988s; Periphery, 258.721 ± 6.367s) (Figure 3F). With respect to the effects of *DNMT* deletion on stress and depression like behavior, PFC DKO mice showed similar corticosterone elevations compared to controls following restraint (Control, n=7, Unstressed, 23.110 ± 10.65ng/mL; Stressed, 173.259 ± 17.142ng/mL; PFC DKO, n=4, Unstressed, 9.012 ± 1.423ng/mL; n=3, Stressed, 176.996 ± 1.306ng/mL) (Figure 3G) and exhibited no differences in forced swim test performance (Control, n=7, Latency, 28.571 ± 7.131s; Immobility, 141.286 ± 16.204s; PFC DKO, n=5, Latency, 53.8 ± 25.781s; Immobility, 138 ± 15.611s) (Figure 3H). Collectively, results from multiple assays suggest that DNMT expression within PFC neurons does not influence affect and the response to stress.

**Figure 3.**
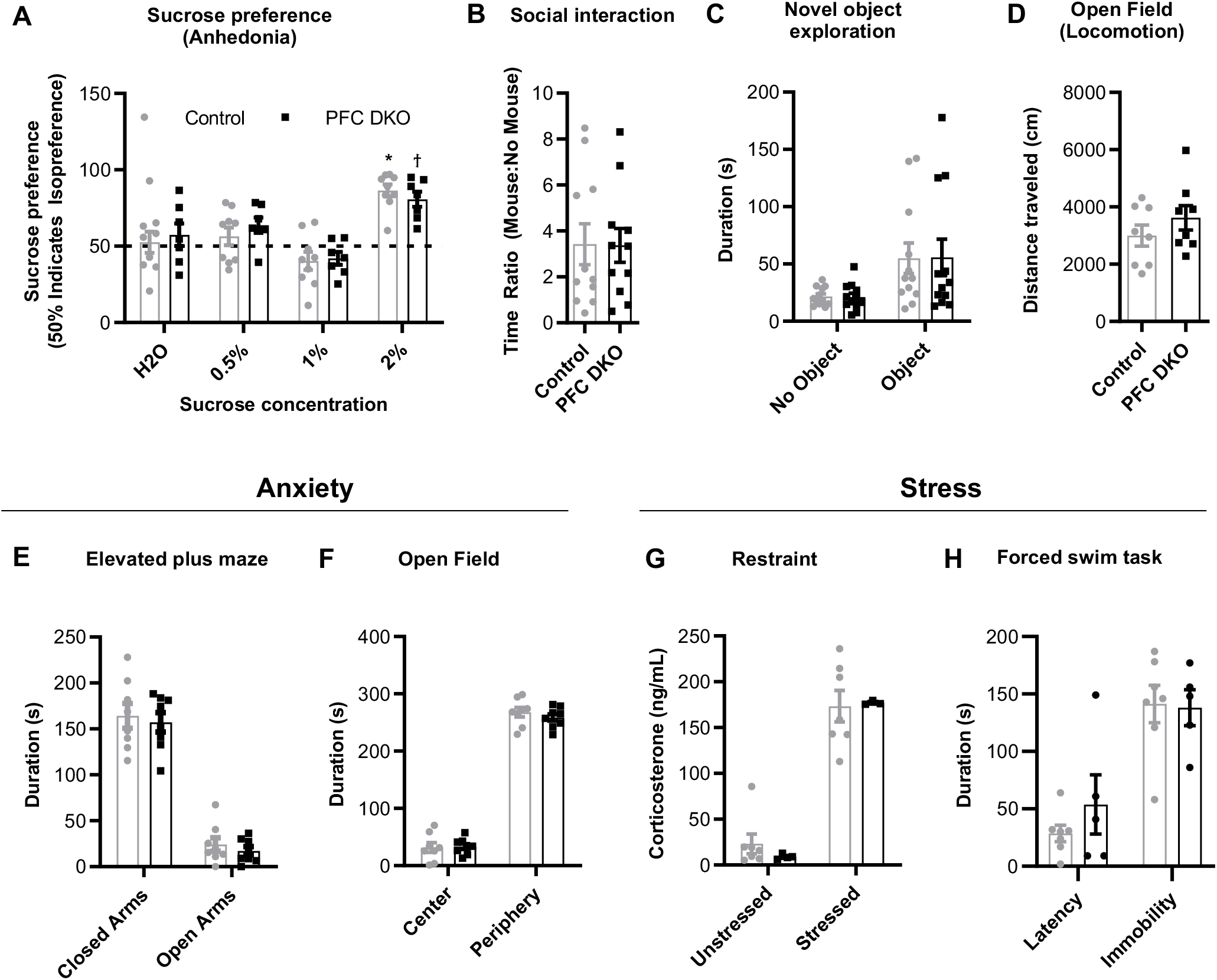
Deletion of *DNMT1* and *DNMT3a* in the mPFC does not alter emotional processing and novelty recognition. A. Preference for sucrose by control and PFC-DKO groups showing no alteration in anhedonia. (control n=9, PFC-DKO n=7 2way-ANOVA multiple comparisons to no sucrose control, \**p*<0.05). B. Ratio of time spent in the zone of interaction with a novel mouse versus time spent investigating the empty restrainer. DKO mice show no difference in social interaction compared to controls. (n=11, *t*-test *p*=0.9664). C. Time spent in the interaction zone exploring a novel object (n=12, *t*-test). D. Total distance travelled by control and PFC-DKO groups while exploring the open field showing no difference in their general locomotion and ability to explore (n=12, *t*-test). E. Time spent in the open and closed arms of an elevated plus maze by control and PFC-DKO groups showing no difference in the anxiety levels between groups (n=8, *t*-test). F. Time spent in the central zone and the periphery while exploring the open field showing no difference in the anxiety levels between groups (n=8, *t*-test). G. Circulating levels of corticosterone in control and PFC-DKO mice showing similar levels of stress reactivity between the groups (control n=7, PFC-DKO: unstressed n=4, stressed n=3, *t*-test). H. Latency to become immobile and duration of immobility during a forced swim task (control n=7, PFC-DKO n=5, *t*-test). Data information: error bars are ± SEM

### PFC deletion of *DNMT1 and DNMT3a* impairs the extinction of conditioned fear behavior

We next investigated whether DNMT expression in the PFC was required for the regulation of PFC-dependent learning. To this end, we determined the effect of *DNMT1* and *DNMT3a* deletion within the PFC on conditioned fear memory extinction, as prior work had shown the requirement of *DNMT1* and *DNMT3a* expression throughout the forebrain in trace fear conditioning (Morris et al., 2014). Mice were conditioned to associate a tone presentation (conditioned stimulus) with foot shock (unconditioned stimulus). Following training, subjects were observed for freezing behavior following tone presentation at 24h, 48h, 72h, and 96h. Interestingly, PFC DKO mice developed abnormal extinction freezing behavior to tone presentation (Figure 4A). While the control mice were able to dissociate the conditioned and unconditioned stimuli (n=7, 24 h, 60.693 ± 8.301%; 48 h, 44.276 ± 5.293%; 72 h, 39.387 ± 7.384%; 96 h, 26.224 ± 3.488%;), PFC DKO (n=7, 24 h, 57.533 ± 5.378%; 48 h, 52.927 ± 8.445%; 72 h, 51.351 ± 7.747%; 96 h, 44.913 ± 9.254%) consistently showed conditioned response even up to 96h post training. Similar deficits in extinction of fear memory in PFC DKO mice was observed in contextual conditioning (Control, n=7, 24 h, 37.94 ± 3.256%; 48 h, 29.856 ± 6.708%; 72 h, 19.643 ± 3.279%; 96 h, 18.317 ± 5.691%; PFC DKO, n=7, 24 h, 47.896 ± 7.827%; 48 h, 37.71 ± 5.237%; 72 h, 48.504 ± 8.96%; 96 h, 45.083 ± 7.89%;) (Figure 4B). Thus, our data was in agreement with prior studies (Morris et al., 2014), further refining the role of action in the PFC of DNMT expression on fear driven learning.

**Figure 4.**
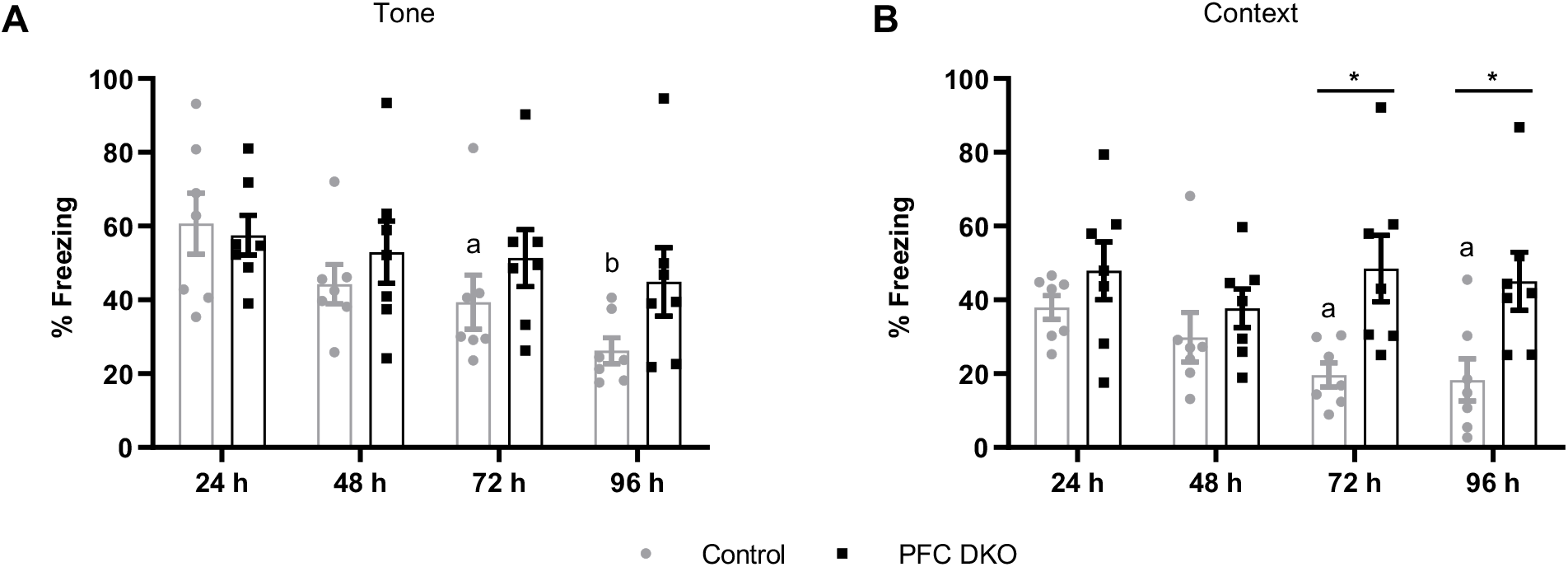
Deletion of *DNMT1* and *DNMT3a* in the mPFC causes abnormal extinction of conditioned fear memory. A. Percent freezing in control and PFC-DKO groups for the presentation of tone (conditioned stimulus) without foot shock (unconditioned stimulus), at indicated time points after the training session. A significant reduction in the %freezing after 72h marks the beginning of extinction of the tone-conditioned fear memory in the control group but not in the PFC-DKO group. (n=7, 2way ANOVA, Dunnett’s multiple comparison test vs 24h a *p*<0.05, b *p*<0.001). B. Percent freezing in control and PFC-DKO groups for the presentation of contextual environment (conditioned stimulus) without foot shock (unconditioned stimulus), at indicated time points after the training session. A significant reduction in the %freezing after 72h marks the beginning of extinction of the context-conditioned fear memory in the control group but not in the PFC-DKO group. (n=7, 2way ANOVA, Dunnett’s multiple comparison test vs 24h a *p*<0.05. 2way ANOVA, Sidak multiple comparison test vs WT \**p*<0.05)

### *DNMT1 and DNMT3a* deletion alters intrinsic membrane properties of PNs in a cortical lamina-specific manner

To begin to determine how loss of the DNMT enzymes could be producing a change in food intake we next sought to investigate how *DNMT* deletion produces functional changes in PNs, the principal output excitatory neurons of PFC. We assessed intrinsic membrane properties and biophysical action potential parameters in both layer-II/III (L-II/III) and layer-V (L-V) PNs (Table 1 and 2). PNs in L-II/III of PFC DKO displayed increased input resistance (*p*=0.0348), reduced rheobase (*p*=0.0348) and action potential amplitude (*p*=0.0114), while also exhibiting a more depolarized action potential threshold (*p*=0.0095) (Figure 5A-D, table 1). Furthermore, the action potential waveform of L-II/III neurons in PFC DKO was found to be altered. L-II/III neurons displayed increased downstroke velocity (*p*=0.0156) and had a negligible change in upstroke velocity (*p*=0.1192) (Figure 5E and table 1). Further, although not statistically significant, action potential durations firstADP50 (*p*=0.0679) and lastADP50 (*p*=0.0555) trended towards being shorter in *Dnmt1* and *Dnmt3a* null neurons (table 1). In addition to this, the firing frequency following current injection trended towards an increase in excitability in the *DNMT* knockouts (Figure 5G and 5H).

**Figure 5:**
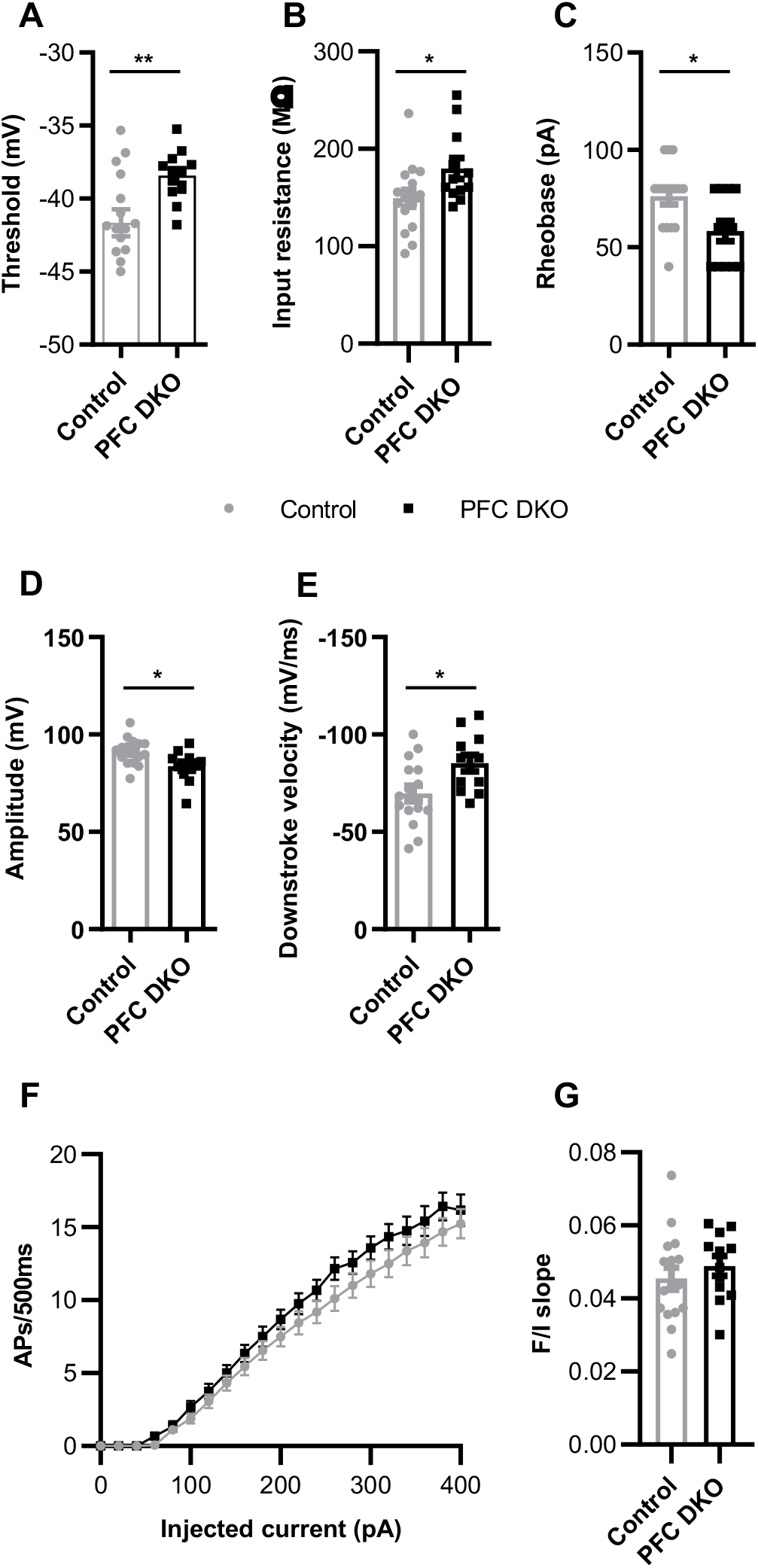
*DNMT1* and *DNMT3a* deletion enhances the excitability of layer-II/III pyramidal neurons in the mPFC. A. Action potential threshold voltage in L-II/III PNs from control and PFC-DKO groups showing a relative increase in DKO mice. (*t*-test, \*\**p*<0.01). B. Input resistance of L-II/III PNs from control and PFC-DKO groups showing a relative increase in DKO mice. (*t*-test, \**p*<0.05). C. Rheobase of L-II/III PNs from control and PFC-DKO groups showing a relative decrease in DKO mice. (Mann-Whitney test, \**p*<0.05). D. Action potential amplitude in L-II/III PNs from control and PFC-DKO groups showing a decrease in DKO mice. (*t*-test, \**p*<0.05). E. Downstroke velocity in L-II/III PNs from control and PFC_DKO groups showing an increase in DKO mice. (*t*-test, \**p*<0.05). F. Frequency-input (FI) current curves for L-II/III PNs from control and PFC-DKO groups showing an increasing trend in the response to a 500ms depolarizing step input current in the DKO animals. (2-way ANOVA with repeated measures, group *p*-0.2132). G. Slopes of linear regression (60-400 pA) of FI curves in D. (*t*-test, *p*=0.4194). Data information: error bars are ± SEM. Number of neurons (n) were from three mice in each group (control n=16, PFC-DKO n=12).

**Table 1:** Comparison of intrinsic electrophysiological membrane properties of layer 2/3 pyramidal neurons.

| Layer II/III | Control<br>(n=16,3 mice) | PFC-DKO<br>(n=12, 3 mice) | p-value |
| --- | --- | --- | --- |
| Resting Membrane Potential (mV) | -70.7 ± 1.6 | -69.5 ± 1.0 | 0.5593 |
| Threshold (mV) | **-41.7 ± 0.9** | **-38.4 ± 0.5** | 0.0095 |
| Upstroke Velocity (mV/ms) | 365 ± 27 | 307 ± 22 | 0.1192 |
| Downstroke Velocity (mV/ms) | **-69.7 ± 4.1** | **-85.3 ± 4.2** | 0.0156 |
| Amplitude (mV) | **91.3 ± 1.7** | **83.8 ± 2.3** | 0.0114 |
| Input Resistance (MΩ) | **149 ± 9** | **180 ± 11** | 0.0348 |
| First APD50 at 400pA (ms) | 1.38 ± 0.09 | 1.13 ± 0.09 | 0.0679 |
| Last APD50 at 400pA (ms) | 2.11 ± 0.17 | 1.67 ± 0.12 | 0.0555 |
| Rheobase (pA) | **76.3 ± 4.2** | **58.3 ± 5.2** | 0.0177 |

**Table 2:** Comparison of intrinsic electrophysiological membrane properties of layer 5 pyramidal neurons. Mean ± SEM are presented and the difference between the group was tested by t-test except Rheobase (Mann-Whitney test). Bolded entries indicate a significant difference between control and DKO mice.

| Layer V | Control<br>(n=13,3 mice) | PFC-DKO<br>(n=14, 3 mice) | p-value |
| --- | --- | --- | --- |
| Resting Membrane Potential (mV) | -69.7 ± 1.6 | -68.9 ± 1.1 | 0.6689 |
| Threshold (mV) | -41.0 ± 1.0 | -39.3 ± 0.7 | 0.1789 |
| Upstroke Velocity (mV/ms) | **371 ± 22** | **275 ± 19** | 0.0035 |
| Downstroke Velocity (mV/ms) | -81.4 ± 3.7 | -71.1 ± 5.2 | 0.1022 |
| Amplitude (mV) | 87.9 ± 2.1 | 87.6 ± 1.4 | 0.9619 |
| Input Resistance (MΩ) | **127 ± 11** | **173 ± 16** | 0.0449 |
| First APD50 at 400pA (ms) | **1.15 ± 0.05** | **1.37 ± 0.06** | 0.017 |
| Last APD50 at 400pA (ms) | **1.58 ± 0.08** | **2.18 ± 0.11** | 0.0004 |
| Rheobase (pA) | 90.8 ± 10.8 | 77.1 ± 13.2 | 0.1818 |

Unlike L-II/III neurons, we did not see the same enhancement in parameters indicative of neuronal excitation in layer V PNs (Table 2). L-V neurons of PFC DKO displayed a significant increase in input resistance (*p*=0.0449) (Figure 6A, table 2) but the concomitant decreasing trend observed in the rheobase was not significant (*p*=0.1818) (Table-2). Interestingly, while upstroke velocity was reduced (*p*=0.0035) (Figure 6B), there was also a trend in the reduction of downstroke velocity (*p*=0.1022; table 2). The action potential durations firstAPD50 (*p*=0.017) (Figure 6C, table 2) and lastAPD50 (*p*=0.0004) (Table 2) were significantly longer in L-V neurons of PFC DKO as compare to those of controls, opposite of what was observed in layer II/III neurons. Finally, the PFC DKO neurons displayed no change in firing frequency in response to depolarizing current injection when compared to controls (Figure 6D and 6E), suggesting that intrinsic excitability of L-V neurons is altered to a lesser extent than that of L-II/III neurons by the absence of DNMT expression.

**Figure 6.**
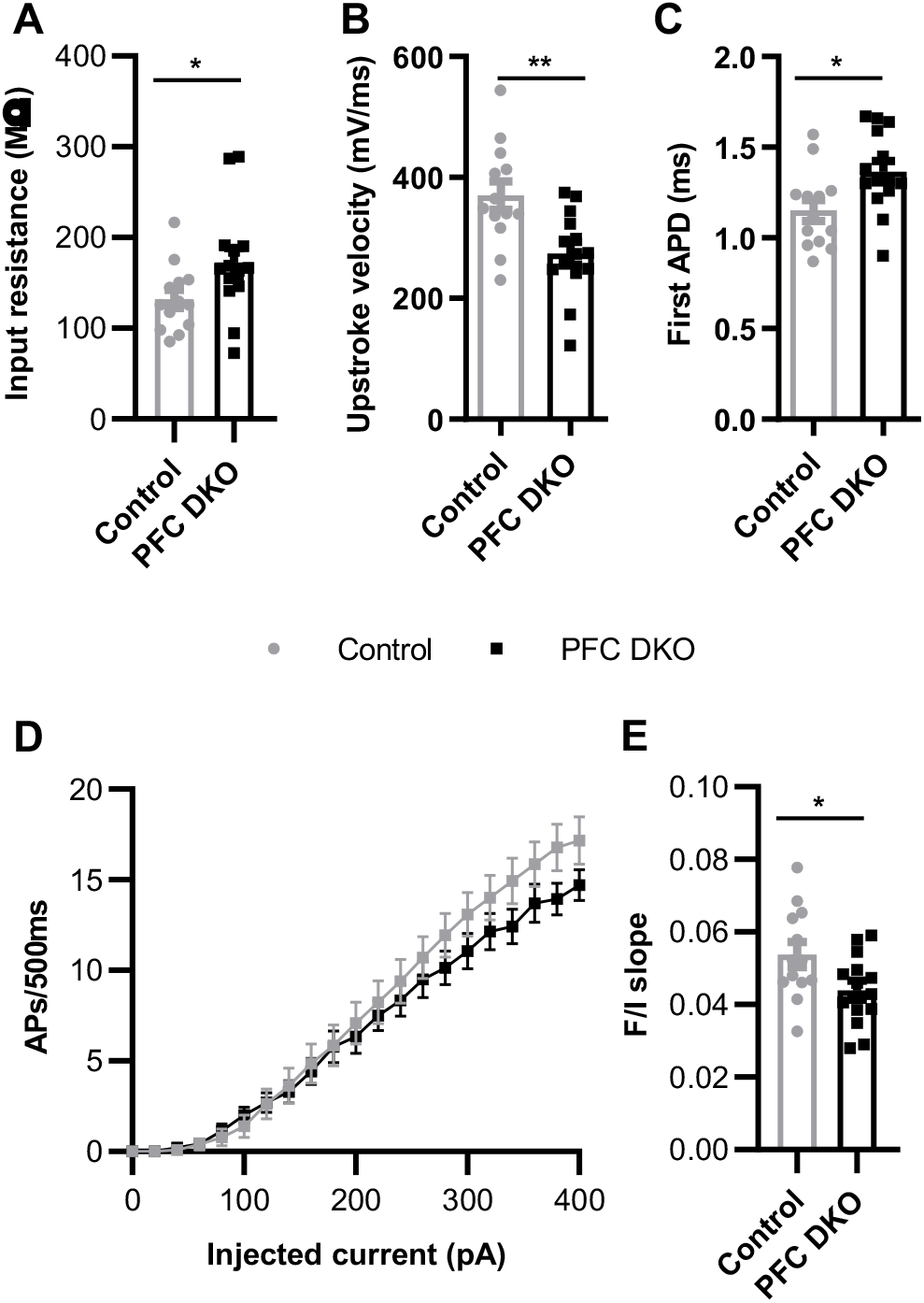
*DNMT1* and *DNMT3a* deletion attenuates the excitability of layer-V pyramidal neurons in the mPFC. A. Input resistance of L-V PNs from control and PFC-DKO groups showing a relative increase in the DKO mice. (*t*-test, \**p*<0.05). B. Upstroke velocity in L-V PNs from control and PFC-DKO groups showing a decrease in the DKO mice. (*t*-test, \*\**p*<0.01). C. Duration of first action potential measured at 50% of amplitude in L-V PNs from control and PFC-DKO groups showing an increase in the DKO mice. (*t*-test, \**p*<0.05). D. Frequency-input (FI) current curves for L-V PNs from control and PFC-DKO groups showing decreasing trend in the response to a 500ms depolarizing step input current in the DKO mice. (2-way ANOVA with repeated measures, group *p*=0.4366). E. Slopes of linear regression (60-400 pA) of FI curves in D. (*t*-test, \**p*<0.05) Data information: error bars are ± SEM. Number of neurons (n) were from three mice in each group (control n=13, PFC-DKO n=14).

### *DNMT1 and DNMT3a* deletion dysregulates DNA methylation selectively at RARβ transcription factor binding sites

DNA methylation has been identified as a crucial factor regulating both Hebbian and homeostatic forms of plasticity. While many of the gene expression changes that result from alterations in DNMT expression have been described, no systematic investigation of the genome wide methylation changes that result from this deletion has been pursued. Thus, to begin to understand how *DNMT* deletion caused the observed behavioral and electrophysiological changes, we profiled the methylomes of the PFC PNs. Using a flow-cytometry-based approach, we were able to purify SATB2 stained and GFP positive neurons form the PFC of both control and PFC DKO animals. We assessed the methylation status of 1,176,292 and 5,671,600 unique CpG and CpH sites with at least 10 reads in one sample from each group using a reduced representation bisulfite sequencing (RRBS-seq) assay. Overall, the majority of CpGs were methylated, while more than 15% of cytosines in the non-CpG context were also significantly methylated (Figure 7A). As expected, the DKO samples showed a significant reduction in the overall methylation of CpG sites when compared to controls. However, the fraction of non-CpG methylated cytosines was higher in the PFC-DKO mice (Figure 7A). In addition to a global decrease in methylation, hypermethylation was also observed at select loci when compared to the control population. With our filtering criteria of at least a 30% change in methylation with *q* < 0.01, we found 71,842 hypermethylated and 76,867 hypomethylated CpG sites in the PFC DKO PN cells (Figure 7B). Similarly, there were 127,483 hypermethylated and 122,263 hypomethylated CpH sites. We next mapped these differentially methylated CpG cites onto 500bp genomic tiles to identify differentially methylated regions. Out of these, we removed 12,774 tiles harboring both hypo and hypermethylated sites (Figure 7B) and used the resulting set of hypo and hyper methylated CpG regions for downstream analysis.

**Figure 7:**
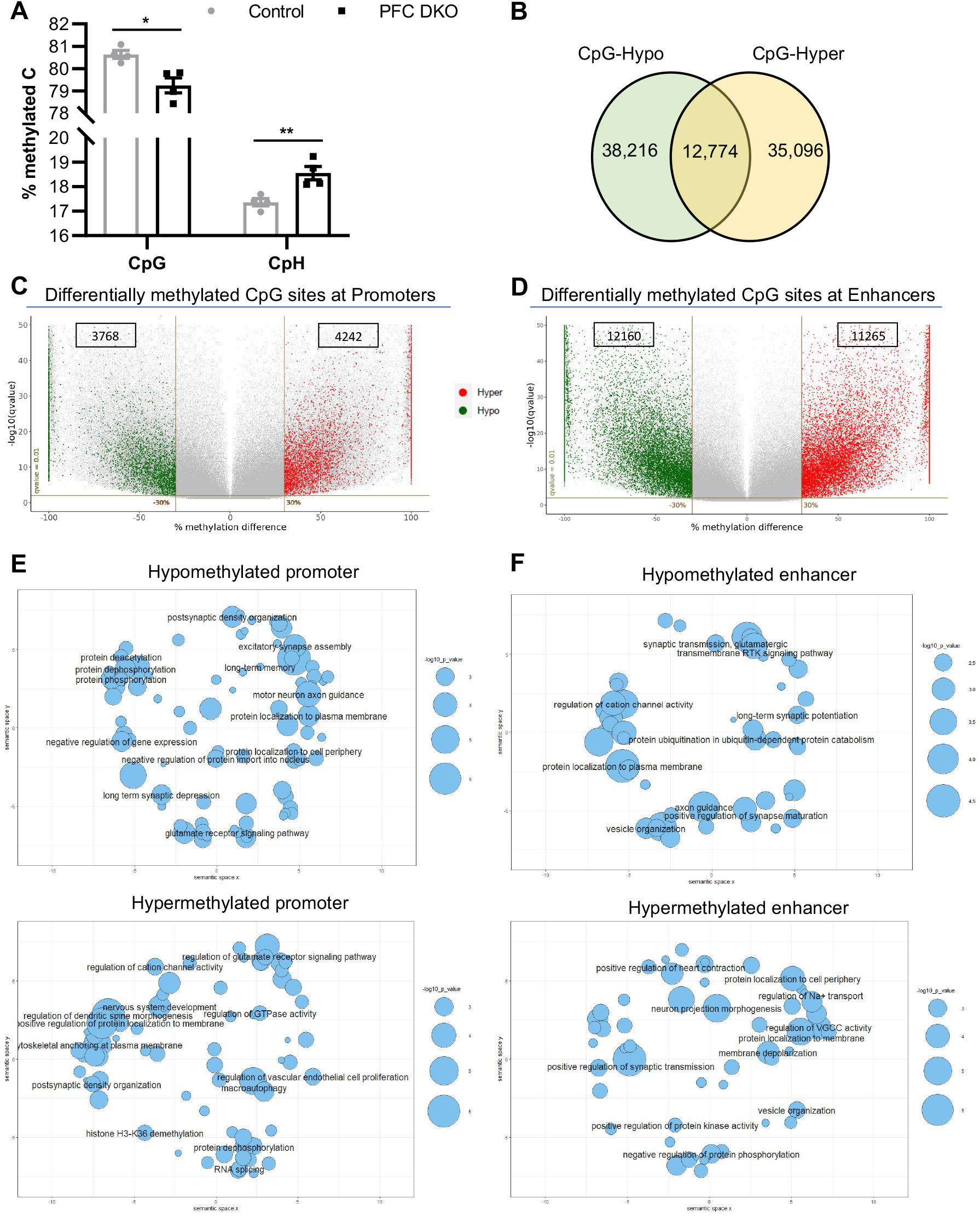
*DNMT1* and *DNMT3A* deletion alters methylation status of regulatory elements of ion channel and synaptic transmission related genes. A. Percentage of methylated cytosines in CpG and CpH (non-CpG) contexts in the control and PFC-DKO groups. Methylation in the CpG context was reduced while that of CpH contest was increased significantly in the PFC-DKO (n=4, *t*-test, \**p*<0.05, \*\**p*<0.01). B. Venn diagram showing hypo and hyper-methylated tiles in PNs of PFC-DKO. The overlapping region contains both hypo and hypermethylated CpG sites which were removed before downstream functional analysis. C. Volcano plot showing the hypo and hypermethylated CpG sites at putative excitatory neuron-specific promoters (GSE63137). Numbers inside the rectangle are the number of differentially methylated CpG sites overlapping with the promoter regions in the group. D. Volcano plot showing the hypo and hypermethylated CpG sites at putative excitatory neuron-specific enhancers (GSE63137). Numbers inside the rectangle are the number of differentially methylated CpG sites overlapping with the promoter regions in the group. E. Gene ontology visualization scatter plots of significantly enriched biological processes (BPs) in hypo (top) and hypermethylated (bottom) promoter-associated genes. F. Gene ontology visualization scatter plots of significantly enriched BPs in hypo (top) and hypermethylated (bottom) enhancer-associated genes. Enrichr (https://maayanlab.coud/Enrichr/) was used to perform functional annotation of differentially methylated regulatory regions. Gene ontology visualization scatter plots were generated in R studio using the codes generated by REVIGO (http://revigo.irb.hr/). Significantly enriched (*p*<0.01) BPs were plotted on a semantic similarity-based scatter plot showing closely related processes as clusters. BPs with least *p*-value or relevance to neurons within each cluster of BPs are labeled. Size of the bubble indicates −log(*p*-value).

Considering the important role of DNA methylation within promoters and enhancers on regulation of transcription, we sought to further characterize whether known gene regulatory regions showed differential DNA methylation following *DNMT* deletion. To this end, we used publicly available datasets to identify active regulatory regions found in cortical PNs using H3K4me3 marked regions to identify promoters and H3K4me1 and H3K27ac dual marked regions to identify enhancers (Mo et al., 2015). Within the data set previously reported by Mo et al (Mo et al. 2015) 13,817 promoters were marked with H3K4me3 and 70,136 enhancer regions marked with H3K4me1 and H3K27ac. Of these, 9,348 regions marked with both promoter and enhancer epigenetic marks and were excluded from the set of enhancers as they were already counted in the promoter set, resulting in the identification of 13,817 promoter and 59,476 enhancer regions in our data set for analysis. Subsequently, we identified 2,940 and 3,087 putative hypo and hypermethylated promoters and 4,821 and 4,421 hypo and hypermethylated putative enhancers, respectively. For our downstream analysis we refer to these regions as differentially methylated promoters (DMPs) and enhancers (DMEs). Analysis of difference in % methylation for CpG sites within these DMPs and DMEs revealed that both complete and partial change in the methylation levels both in the case of promoters (Figure 7C) and enhancers (Figure 7D). Interestingly, several known molecular regulators of feeding behavior were found to have differentially methylated promoter or enhancer regions. For example, all three opioid receptors *OPRD1*, *OPRK1*, and *OPRM1* were differentially methylated. While the distal enhancer of *OPRK1* and *OPRM1* was hypermethylated, multiple promoters and enhancers of *OPRD1* were hypo and hypermethylated. Furthermore, regions associated with both endogenous opioids were differentially methylated. The promoter of proenkephalin (*PENK*) was found to be hypermethylated with promoters and enhancers associated with prodynorphin (*PDYN*) being hypomethylated. With respect to melanocortin signaling, distal enhancers of the melanocortin receptors *MC1R*, *MC2R* and *MC3R* were found to be hypermethylated. Further, an additional proximal enhancer region of *MC3R* was hypomethylated although no differentially methylated regions were associated with *MC4R*. Among the receptors for orexin, *HCRTR1* regulatory regions were found to be hypo and hypermethylated with the *HCRTR2* promoter found to be hypomethylated. Multiple enhancers regulating the expression of the ghrelin receptor (*GHSR*) were differentially methylated while multiple promoters and enhancer regions associated with pro-opiomelanocortin (POMC), and the processing enzymes that cleave this transcript such as Carboxypeptidase E (*CPE*), Peptidyl-glycine α-amidating monooxygenase (Pam), Proprotein convertase subtilisin/Kexin Type 1 (PCSK1) were differentially methylated. Thus, our analysis suggests that many genes associated with the regulation of food intake that show expression in the frontal cortex show methylation patterns that are regulated by DNMT expression, suggesting the potential for transcription dysregulation that could lead to an alteration in the drive to consume palatable food.

We then investigated whether more general processes known to be involved in controlling neuronal excitability and signal transduction were affected by *DNMT* deletion. Functional analysis of genes associated with DMPs and DMEs identified several key biological processes (BP) linked to the control of neuron function, such as signal transmission, ion channel activity regulation and protein localization to the membrane (Figure 7E and F). Some of the BPs identified, including long term potentiation, depression, synaptic plasticity, and memory, had already been shown to be influenced by DNA methylation in other brain regions (Feng et al., 2010; LaPlant et al., 2010; Levenson et al., 2006; Miller et al., 2008; Morris et al., 2014).

We then investigated the transcription factors that could potentially show functional alterations in response to differential methylation within the identified promoter and enhancer regions. ChEA-analysis from enrichr revealed that a significant fraction of both hypo and hyper methylated regulatory regions harbor RARβ binding sites (Figure 8A), based on data obtained from RARβ transcription factor binding studies conducted in the striatum (Niewiadomska-Cimicka et al., 2017). Of these potential RARB binding sites, our analysis revealed that 776 were hypomethylated while 880 sites were hypermethylated in PFC DKO PN neurons compared to control (Figure 8B). Interestingly, a majority of the differentially methylated RARβ sites were located within 5kb from the TSS of associated genes (Figure 8C and 8E) enriched within a distinct set of biological processes. Hypomethylated-RARβ binding sites (hypo-RARβ) were enriched within genes associated with the regulation of GTPase activity, cation channel activity, vesical mediated transport, and synapse maturation (Figure 8D). Furthermore, hypo-RARβ sites were associated with a host of genes that encoded RAB-family GTPases (*RAB10*, *RAB11B*, *RAB40B* and *RAB40C*), Neuroligin1, Vesicle associated membrane protein 2, Rectifier voltage-gated potassium channels (*KCNB1* and *KCNB2*) and kinases such as calmodulin-dependent protein kinase-2 subunit beta (*CAMK2B*) and neurotrophic tyrosine receptor kinase-2 (*NTRK2*). Interestingly, *RARβ*, *HDAC2* and *TET2* were also in the list of hypo-RARβ-associated genes. On the other hand, regulation of membrane targeted protein trafficking and transcription from the RNA polymerase II promoter were some of the enriched biological processes associated with hypermethylated-RARβ binding (hyper-RARβ) sites (Figure 8F). Furthermore, hyper-RARβ sites were associated with transcription factors such as *CITED2*, *CRCT2*, *NR3C1*, *RARG*, *RORA*, and *JUNB*, and kinases such as *GSK3B*, *AKT3*, *MAP3K1*, *ARF4* and *JAK1*. A complete list of the gene-region association analysis is provided as supplementary information (Supplementary file-2).

**Figure 8:**
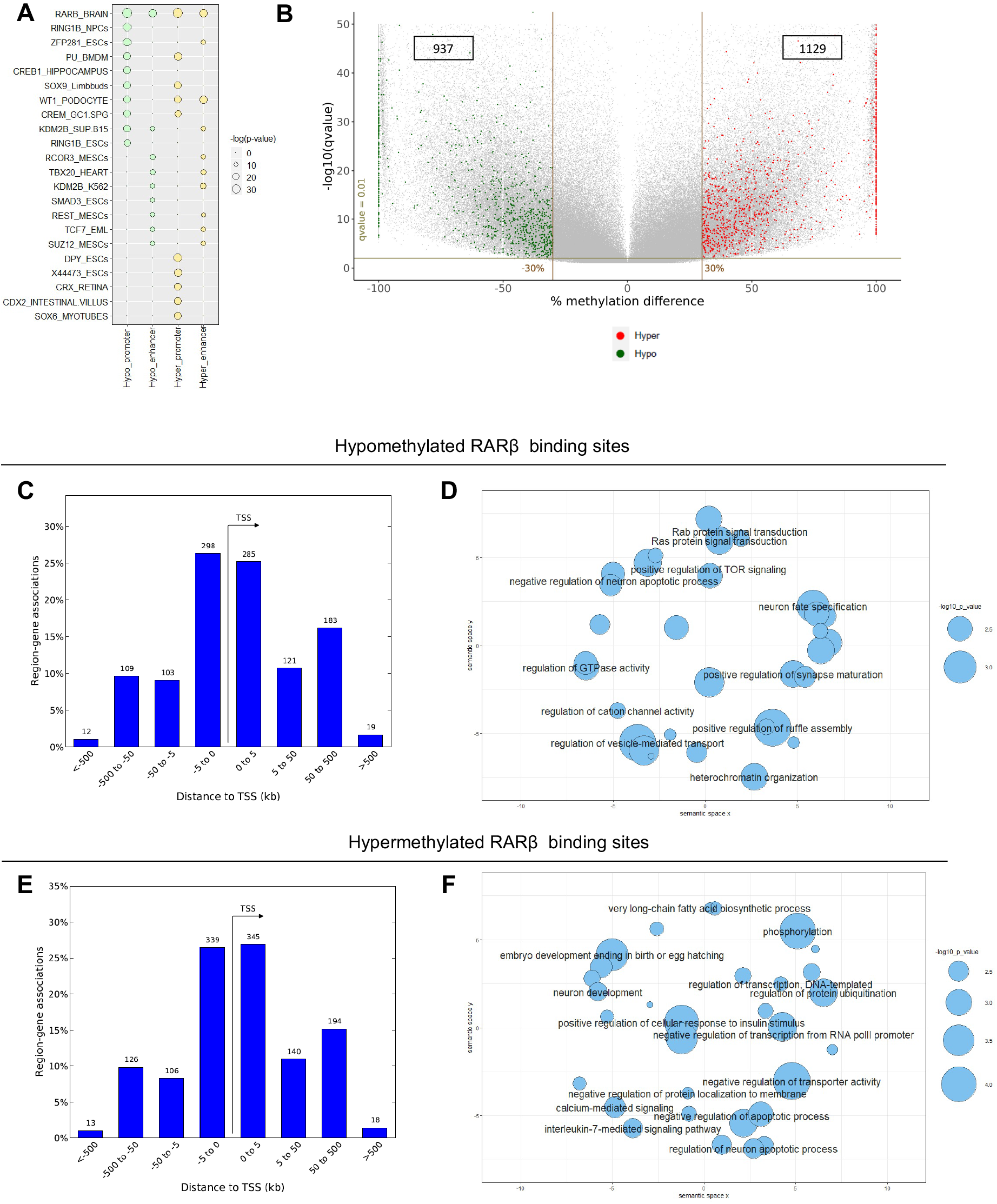
Crosstalk between retinoic acid signaling and DNA methylation in pyramidal neurons. A. Bubble plot showing the top ten enriched regions for various chromatin binding proteins in hypo and hyper methylated promoters and enhancers. Row names are the chromatin-X factor and the associated biosample separated by an underscore. Enrichments were identified by ChEA analysis from Enrichr (https://maayanlab.coud/Enrichr/). Bubble sizes are proportional to −log(*p*-value). B. Volcano plot showing the hypo and hypermethylated CpG sites at RARβ binding sites (GSE67829). Numbers inside the rectangle are the number of differentially methylated CpG sites overlapping with the promoter regions in the group. C. Region-gene association graph showing the distribution of hypomethylated RARβ binding sites around the associated transcription start site (TSS). D. Gene ontology visualization scatter plots of significantly enriched BPs in hypomethylated RARβ binding site-associated genes. E. Region-gene association graph showing the distribution of hypermethylated RARβ binding sites around the associated transcription start site (TSS). F. Gene ontology visualization scatter plots of significantly enriched BPs in hypermethylated RARβ binding site-associated genes. Enrichr (https://maayanlab.coud/Enrichr/) was used to perform functional annotation of differentially methylated regulatory regions. Region-gene association graphs were generated by GREAT (http://great.stanford.edu/public/html/). Gene ontology visualization scatter plots were generated in R studio using the codes generated by REVIGO (http://revigo.irb.hr/). Significantly enriched (*p*<0.01) BPs were plotted on a semantic similarity-based scatter plot showing closely related processes as clusters. BPs with least *p*-value or greatest relevance to neuron function within each cluster of BPs are labeled. Size of the bubble indicates −log(*p*-value).

## Discussion

Our *DNMT* conditional knockout studies have demonstrated the important role of these proteins in the frontal cortical control of animal behavior, neurophysiology, and the methylation status of select loci in the genome.

Deletion of *Dnmt1* and *Dnmt3a* from mouse medial prefrontal cortex resulted in a selective effect on animal ingestive behavior, reducing palatable food intake. Interestingly, while palatable food intake was suppressed, chow food intake was not affected. Furthermore, no effect on chow intake was observed in fasted animals, suggesting that behavioral change in response to our manipulation did not depend as much on the valence of the food consumed, but on the nutrient content. While control and *DNMT* knockout mice showed a similar ability to perceive sucrose, and thus palatability, our manipulation appears instead to have produced a select effect on the amount of palatable food consumed. Furthermore, our RRBS study revealed promoters and enhancer regions associated with several of the known molecular regulators of feeding behavior, including orexigenic (Orexin, Endocannabinoids, Galanin, Melanin-concentrating hormone, endogenous opioids) and anorexigenic agents (POMC and enzymes involved in the processing of this peptide) and their receptors, were found to be differentially methylated. Interestingly, differential methylation of multiple regulatory regions associated with these genes indicates the possible role for DNA methylation in the regulation of cellular state-specific expression of transcript variants.

In support of our observations, prior studies have shown how the frontal cortex could act to either enhance or decrease the consumption of palatable food, depending on the frontal cortex subregion targeted. Berridge and colleagues noted areas of the frontal cortex that could drive, following μ-Opioid receptor agonism, either a suppression or an increase in food intake (Castro and Berridge, 2017). Cottone and colleagues showed how naltrexone could, when applied focally to the frontal cortex, selectively limit palatable food intake (Blasio et al., 2014). Although our targeting of the *DNMT* deletion was not frontal cortex subregion specific, we were still able to produce a select effect on palatable feeding. Future work will, therefore, focus on the identification of the prefrontal cortical subregions and neuronal subtypes responsible for the effect, while also determining how opioid signaling and DNA methylation changes could both produce a phenotypically similar suppression of food intake.

Our studies examining how *DNMT1* and *3A* deletion could dysregulate other behaviors controlled by the frontal cortex demonstrated that a change in anxiety, depression-like behavior, novelty detection or a change in stress responsiveness did not result from our approach. Importantly, these data also suggest that a lack of change in these behavioral parameters confirms the specificity of the observed effect of *DNMT* deletion on food intake.

Interestingly, in disagreement with our data, prior studies did observe a change in affect following manipulation of *DNMT* expression. *DNMT1* deletion throughout the forebrain showed anxiolytic and antidepressant effects, with selective deletion of Dnmt3a producing no effect (Morris et al., 2016). *DNMT3A* deletion, meanwhile, impaired spatial and novel object recognition (Morris et al., 2014), while we observed no change in these behaviors.

When comparing our results to the only other study to examine *DNMT* function selectively in the PFC using a knockdown approach, differences in the effect of *DNMT* deletion on behavior were also noted, with *DNMT3A* knockdown being anxiogenic and *DNMT1* knockdown having no effect on anxiety behavior in this prior work.(Elliott et al., 2016).

Finally, with respect to effects on learning and memory, our data did recapitulate the observed effect of *DNMT1* and *DNMT3A* loss reported in prior work. While we showed how extinction of fear memory was impaired during both a cued and contextual recall, similar results were observed by Morris et al., following the selective deletion of *DNMT3A* throughout forebrain excitatory neurons (Morris et al., 2014).

It is possible that the disagreement of our data with certain prior experiments could be explained by the different methodologies used and cell populations targeted in the analysis of *DNMT* function. For example, unlike our neuron-specific approach, lentiviral-mediated silencing of the DNMT enzymes target both neuronal and non-neuronal cells within the mPFC. This potentially indicates that DNMT expression in the non-neuronal cells of mPFC are crucial in regulating anxiety like behavior. It is also likely that our targeting of excitatory and inhibitory neurons produced an effect on circuit activity that differed significantly from those studies employing a CAMKII-driven Cre recombinase to delete DNMT expression. As these conflicting results show, a future investigation of the cell subtype selective effects of *DNMT* deletion on behavior is warranted.

Our observed effects of *DNMT* deletion on food intake and fear learning resulted from a complex and cortical layer specific effect on intrinsic neuron excitability. L-II/III neurons showed reduced rheobase and increased input resistance indicating pro-excitatory changes, an effect that was also observed in the F-I plots. Similarly, changes in upstroke and downstroke velocities, amplitude, and action potential duration (APD50), all measures of action potential waveform, showed that L-II/III PFC DKO neurons fire smaller and faster action potentials compared to controls due to enhanced repolarization. Interestingly, action potential threshold was also increased, suggesting that, while a greater degree of depolarization was required to initiate an action potential in the L-II/III PFC DKO neurons, these cells were likely to show an elevated firing frequency when compared to controls.

In the case of L-V neurons, the overall effect on neuronal excitability following *DNMT* deletion was significantly different, when compared to the L-II/III neurons. With a reduction in the depolarization rate relative to control neurons, action potentials were longer in the L-V neurons of PFC DKO which would suggest a reduction in evoked firing frequency. However, increase in input resistance suggested an increase in excitability.

Collectively, our electrophysiological analysis of both superficial and deep layer neurons suggests the importance of DNMT expression in the regulation of ion channels, likely at the level of ion channel expression, based on the cell autonomous changes observed in excitability. Our results also suggest that DNMT expression might be crucial in maintaining the layer-specific cellular identity of PNs and that variation in methylation could be involved in establishing the considerable functional and morphological heterogeneity observed within PNs of the PFC (Barthó et al., 2004; Dégenètais et al., 2002; Molnár and Cheung, 2006; Radnikow and Feldmeyer, 2018; Yang et al., 1996). Our electrophysiology experiments did not investigate synaptic changes and therefore further investigation into how the DNMT enzymes change synaptic (as opposed to intrinsic) excitability is warranted to gain further insight into how the deletion of *DNMT1* and *DNMT3A* might influence neuronal physiology and ultimately alter behavior.

With respect to how *DNMT* deletion acts at the level of the genome to produce a change in neuron excitability and subsequently animal behavior, we observed significant bidirectional plasticity at both CpG and CpH residues within a defined set of loci that likely lead to selective effects on neuron function. The percent methylation of a given cytosine within a dynamically methylated region results from the combined activities of both methylation mark writers and erasers (Ginno et al., 2020). Therefore, ablating either methylation writers or erasers would be expected to perturb the methylation kinetics at dynamically methylated regions. Our RRBS results identified such regions within a genetically defined pyramidal neuron population, that could explain both the observed changes in behavior and neurphysiology. Interestingly, we identified a comparable number of hypo and hyper methylated regions in the PFC DKO neurons compared to controls. Considering the demonstrated influence of DNMT enzymes on demethylation and interdependencies in recruitment of DNMT and TET enzymes (Chatterjee et al., 2018; Chen et al., 2013a; Gu et al., 2018; Kangaspeska et al., 2008; Métivier et al., 2008; van der Wijst et al., 2015; Zhang et al., 2017), hyper methylated sites in the PFC DKO neurons are not surprising. Plausibly, this indicates that the presence of DNMT enzymes is essential for demethylation at these hypermethylated sites. Functional annotation of the identified DMRs, meanwhile, revealed a sizable number of excitatory neuron-specific active promoters and enhancers demonstrating the interplay between transcription and dynamic DNA methylation. Furthermore, regulatory regions of genes which code for ion channels and their modulatory subunits were shown to be differentially methylated, likely leading to differential transcription, as demonstrated in prior studies (Meadows et al., 2016).

The impact on both upstroke and downstroke velocities noted in our waveform analysis suggests that both Na+ and K+ channels are regulated by DNA methylation. In line with this, our RRBS data shows the altered methylation status of a variety of ion channels and their modulatory subunits (Supplementary file 2). Interestingly, several DMR-genes were associated with distinct hypo and hypermethylated regions possibly indicating the importance of DNA methylation on the regulation of transcript variants and utilization of alternate promoters, depending on the physiological context. One immediate possibility could also be that these changes might result in the differential expression of transcript variants that code for these and other ion channels. One such example is seen in the CACNB subunit genes that encode the auxiliary b subunits of high voltage activated Ca^2+^ channels, forming the modulatory region at the cytosolic face of the channel.

Retinoic acid (RA) signaling had previously been implicated in the regulation of synaptic activity mostly via RARα-mediated non-transcriptional mechanisms (Chen et al., 2014; Hsu et al., 2019; Maghsoodi et al., 2008; Zhong et al., 2018). However, the genomic actions of RA-signaling on neuronal function are also likely to be significant (Niewiadomska-Cimicka et al., 2017; Nomoto et al., 2012). It is evident from a variety of organ systems that genomic actions of nuclear hormone signaling are closely associated with DNA methylation suggesting nuclear hormone-mediated dynamic DNA methylation could significantly impinge upon transcriptional regulation (Dhiman et al., 2015; Kouzmenko et al., 2010; Sandoval-Hernández et al., 2016).

RARs are known to heterodimerize with RXR and are also shown to recruit DNA erasers, histone acetyl transferases and deacetylases to the regulatory regions (Minucci et al., 1997). Moreover, RARβ transcription shows autoregulation, in addition to promoter DNA methylation at this loci (de Thé et al., 1990; Hayashi et al., 2001). Furthermore, the interconnection between DNA methylation and RA-signaling has been recently described in a non-neuronal system (Hassan et al., 2017). Interestingly, a recent case-controlled clinical study has shown how differential DNA methylation of RARβ is a key factor explaining the association between the selection of healthy lifestyle choices and a reduced risk of breast cancer development (Wang et al., 2020). In light of this reported evidence, our finding that RARβ binding sites were significantly enriched in differentially methylated regulatory regions reinforces the idea that genomic actions of RA signaling could be one of the crucial pathways regulating PFC neuronal activity and associated behavior.

In conclusion, our studies have highlighted the role of two important enzymes that regulate the DNA methylation status of the neuronal genome within the PFC. Deletion of *DNMT1* and *DNMT3a* from neurons of the PFC produced a specific effect on palatable food intake and fear memory, in the absence of any change in affect. Our data reveal a novel effect of the deletion on neuronal physiology, with superficial cortical layer PNs showing enhanced excitation while deep layer PNs exhibited reduced excitability. Finally, our analysis of DNA methylation implicates RA signaling in the control of both neuron physiology and behavior.

## Materials and Methods

### Animals

All experiments were performed in accordance with Association for Assessment of Laboratory Animal Care policies and approved by the University of Virginia Animal Care and Use Committee. Generation of Dnmt1^2lox/2lox^ Dnmt3a^2lox/2lox^ mice has been described previously (Feng et al., 2010). Mice were group housed (up to 5 mice per cage) under standard climate-controlled conditions on a 12 h light-dark cycle (lights on at 6:00 AM) with standard rodent chow (Harlan Teklad, TD7912) and water available *ad libitum*. Six-8 weeks-old male mice were used for all experiments.

### Viral vectors and Stereotaxic injections

For neuron-specific double conditional knockouts of DNMT1 and DNMT3a in mPFC (PFC DKO), we used AAV2.1-Syn-Cre-GFP (UNC vector core) to produce the deletion or AAV2.1-Syn-GFP as a control. Wildtype mice (C57BL6J) injected with same vector served control. Mice were anesthetized with a mixture of ketamine/dexmedetomidine (40 mg/kg / 0.4 mg/kg) and maintained by inhaled isoflurane (1%) during the procedure. Virus (200 nL/side) was delivered to the mPFC bilaterally using coordinates +2.05 mm anterior from Bregma, +/-0.35 mm lateral to midline, and −1.5 mm ventral from dura. Immediately following surgery, mice received subcutaneous injections of antisedan (0.4 mg/kg) and ketoprofen (0.1 mL of 1 mg/mL). Mice were observed 5 days post-surgery and received ketoprofen subcutaneously as needed.

### Acute brain slice preparation and Electrophysiological recordings

Acute brain slice preparation and electrophysiological recordings were performed as described previously (Wengert et al., 2019). Only GFP expressing neurons were patched.

### FACS sorted isolation of frontal cortical nuclei

Mice were anaesthetized with 0.12 mL euthanasia solution (mix information) i.p., immediately followed by cervical dislocation and decapitation. Unfixed PFC tissue was roughly harvested, snap frozen in liquid nitrogen, and stored at −70°C until further processing. Batches of tissue punches were processed for nuclear isolation as described previously (Grindberg et al., 2013). Briefly, individual tissue punches were incubated in a Tris-buffered magnesium and potassium salt solution containing protease inhibitor and dithiothreitol. Tissue was triturated using a 1 ml pipette and was then homogenized using a polytron homogenizer. Triton-X100 was then added and the tissue Dounce homogenized using two pestles of decreasing size. Homogenate was strained through a 70 μm filter and centrifuged at 1000 x g for 8 min at 4°C. Nuclei pellet was resuspended, layered on a 29% Iodixanol gradient and centrifuged at 10,300 x g for 20 min at 4°C. Supernatant was collected, nuclei were counted from an aliquot. Nuclei were stained with propidium iodide (PI) before sorting. Both PI-positive and double positive (PI and GFP) nuclei were FACS-sorted.

To isolate nuclei pyramidal neurons of the mPFC, nuclei in supernatant were incubated for 1 h with DAPI and anti-Satb2 antibody (1:100, abcam ab34735) conjugated to R-Phycoerythrin (RPE) using the Lightening-Link^®^ R-Phycoerythrin conjugating kit (703-0010, Innova Biosciences), performed as per manufacturer’s instructions. Double positive (RPE and DAPI) and triple positive (GFP, RPE and DAPI) nuclei were FACS-sorted. FACS sorting was performed using a BD FACSVantage SE Turbo Sorter with FACSDiVa Option (Flow Cytometry Core Facility, University of Virginia). FACS-sorted nuclei were spun down at 500 x g for 10 min at 4°C, nuclei pellet was snap frozen in liquid nitrogen and stored at −70°C until further processing.

### Real-time PCR

cDNA was synthesized from the FACS-sorted nuclei using the TaqMan^®^ Gene Expression Cells to CT kit (ThermoFisher) as per manufacturer’s instructions. TaqMan^®^ assays for Dnmt1 (Mm01151063_m1) and Dnmt3a (Mm00432881_m1) were used to quantify the relative expression levels. TaqMan^®^ assay for 18s (Hs99999901_s1) served as internal control. Fold mRNA expression was calculated using the delta-delta CT method. All PCR reactions were performed using the MyIQ Single Color Real-Time PCR detection system (Bio-Rad).

### RRBS library preparation and sequencing

Reduced Representation Bisulfite Sequencing libraries were prepared as described previously (Bock et al., 2010).

### RRBS data analysis

Quality of sequencing reads was checked by Fastqc. Reads were trimmed for low-quality bases (phred score < 30) and adaptor sequences by Trimmomatic-0.32. Reads shorter than 18 bp were discarded. Bismark program (version 0.19.1), in combination with Bowtie2, was used to map the reads to mouse genome (mm10) and subsequent methylation calling. CX-report containing methylation calls for all Cytosine contexts was generated and for downstream analysis CpG and CpH contexts were separated. methylKit (version 0.99.2) identified the differentially methylated CpGs (dmCpGs). dmCpGs were mapped on to 500bp genomic tails to identify hypo and hypermethylated genomic regions. Genomic regions with hypo and hypermethylated CpGs were excluded from the analysis. All the arithmetic operations on genomic intervals were performed by bedtools. For excitatory neuron-specific enhancers and promoters, GSE63137 dataset was used. As described by Mo et al., genomic regions marked with H3K4me3 were considered as promoters and those regions marked with H3K4me1 and H3K27ac were considered as enhancers (Mo et al., 2015). Rarβ binding sites were from dataset GSE67829. Region-gene association, ChIP-X Enrichment Analysis (ChEA) and functional annotation of associated genes was done by Enrichr (Chen et al., 2013b). REVIGO web tool was used to plot enriched biological processes as semantic similarity-based scatter plots (Supek et al., 2011). Codes from REVIGO were processed in R. Wherein similar biological processes were clustered together and processes with most significantly enriched or closely related to neuronal function within a cluster was labeled on the plot. Complete lists of gene ontology analysis are provided as supplementary information (Supplementary file 1).

### Brain tissue processing and fluorescence microscopy

Brain slice preparation was performed exactly as described previously (Gaykema et al., 2014). PFC slices were mounted with a VECTASHIELD antifade mounting medium with DAPI (H-1200-10, Vector Laboratories). Complete slice was tile scanned under a 10× objective of Olympus upright fluorescence microscope. Images were stitched by Neurolucida software.

### Behavior Experiments

### Binge-like feeding

Mice were individually housed on sani-chip bedding in the presence of a 100 mm petri dish bottom and allowed to habituate for 3 days prior to the feeding assay. One day prior to testing, subjects were offered a small (~0.25g) sample of either a new piece of chow, high fat, high sugar diet (HFD; Research Diets Inc., #D12331, 5.56 kcal/g), or Western Diet (WD; Envigo, #TD88137, 4.41 kcal/g) on the petri dish. At lights on the next day, a new full pellet of regular chow or HFD or WD was placed in the petri dish and subjects were allowed to free feed for 60 min. The binge feeding assay was repeated for 2 subsequent days.

### Daily food intake

Mice were individually housed, and food consumption was monitored daily for 5 days.

### Social interaction

Mice were allowed to explore a 41”x41” open square arena with a perforated acrylic cell located in the center of one of the arena walls for 2.5 minutes. Immediately following this session, a novel mouse was placed in the perforated cell and subjects were once again allowed to explore the arena for 2.5 minutes. Subjects always entered the arena in the far corner opposite of the perforated cell. EthoVision tracking software (Noldus) was used to capture mouse position within the arena. The arena was subdivided into the following regions: interaction zone, periphery, and corners and data presented as a ratio of time spent within the interaction zone with and without the novel mouse.

### Sucrose preference

Subjects were given free access to two sipper tubes, one containing sucrose and the other water, for 14 days. Sipper tube position was alternated daily. For the first 3 days, both tubes were filled with water. Subjects were then exposed to increasing concentrations of sucrose solutions starting from 0.5% (days 4-6), 1.0% (days 8-10) to 2.0% (days 12-14). On days 7 and 11, both sipper tubes were filled with water.

### Elevated plus maze

Subjects were allowed to explore an elevated plus maze (open arms and closed arms were both 24” in length) for 5 min. Mouse location was tracked using the EthoVision tracking system. Time in open and closed arm was measured.

### Open field

Subjects were allowed to explore a 41”x41” open square arena for 5 min. Time in center and periphery, and total distance travelled were measured.

### Novel object recognition task

Mice were allowed to explore a 41”x41” open square arena for 2.5 minutes. Immediately following this session, a novel object (5cm copper pipe elbow joint) was placed in the middle of the arena and subjects were once again allowed to explore the arena for 2.5 minutes. Subjects always entered the arena in the same corner of the arena. EthoVision tracking software (Noldus) was used to capture mouse position within the arena. The arena was subdivided into the following regions: interaction zone, periphery, and corners and data presented as time spent within the interaction zone with and without object.

### Blood corticosterone measurements

For baseline corticosterone measurements, subjects were habituated for 3 days to the restraint device and tail massage motion. On the 4th day, whole blood samples were collected via tail vein in non-heparin glass capillary tubes and transferred to microcentrifuge tubes during the late afternoon (4pm-5pm). On the 5th day, whole blood samples were collected again during the morning (7am-8am) via tail vein. For corticosterone measurements under acute stress, mice were placed in restraints fashioned from modified 50 mL conical tubes for 30 min and blood sample was collected. There was a two-week refractory period between baseline and acute stress measurements. Blood serum was isolated from whole blood samples by incubating samples at room temperature for 10 min followed by centrifugation at 2000×g for 10 min at 4°C. Serum in the supernatant was stored at −20°C for further analysis. Corticosterone was assayed using the Corticosterone rat/mouse ELISA (Immuno-Biological Laboratories Inc., #IB79175) following the manufacturer’s protocol.

### Statistical analyses for behavioral data

Unpaired t-test compared the differences between two groups. Multiple groups were compared by ANOVA. Data are presented as mean ± SEM (represented as error bars). *p*-value<0.05 was considered statistically significant.

